# Recycling of prefrontal subspaces dynamically multiplexes information

**DOI:** 10.1101/2025.05.02.651965

**Authors:** Pablo T. Wentz, Scott L. Brincat, Anitha Pasupathy, Earl K. Miller

**Author notes:** These authors contributed equally (Pablo T. Wentz, Scott L. Brincat).

## Abstract

Cortical spiking activity in areas like prefrontal cortex (PFC) encodes information only along a small number of dimensions, a subspace of the full space of population activity patterns. PFC often uses distinct subspaces to represent incoming sensory information and to maintain it in working memory. However, it’s generally assumed that a subspace carries specific information within a specific time frame. Here, in two datasets, we show that neither assumption is correct. The same subspace used during initial encoding was later reactivated during the behavioral response. We also found that subspaces ostensibly dedicated to processing sensory information were recycled to represent behavioral response information later in a trial. We further found a consistent spatial mapping between the original subspace usage and its later recycling. These results demonstrate previously unknown flexibility of PFC subspace coding, both in the deployment of subspaces across time and in their multiplexing of information. This flexibility suggests that PFC subspaces function more like “workspaces” into which different types of information can be written.

## Introduction

Cognition is fundamentally flexible. The brain must constantly integrate multiple streams of information with internal goals to generate decisions and actions. Because internal goals frequently change, this process requires commensurate flexibility in sensory-motor mappings. We sought to examine how neural populations are configured and reconfigured to meet changing task demands.

The prefrontal cortex (PFC) is particularly critical for these processes. It receives bottom-up signals from all sensory systems and top-down signals reflecting current goals and other internal processes^1^. Prefrontal processing is thought to have a particularly high degree of flexibility that facilitates diverse input/output mappings^2–4^. Rather than exclusively coding for one aspect of a cognitive task, single PFC neurons often multiplex different types of information^3,5^.

At the level of single neurons, this multiplexing introduces ambiguity: the same neuron’s activity can represent different meanings depending on context. This ambiguity is thought to resolve when considering the pattern of activity across a neural population^6–8^. Recent studies have shown that high-dimensional population activity is highly coordinated across neurons. Instead of using the full spectrum of possible spiking patterns, neural populations organize information along “subspaces”, low-dimensional subsets of activity. Structuring activity in such subspaces is believed to flexibly organize information processing^9–11^ and communication^12^.

Studies have shown that PFC population coding is dynamic. Distinct activity subspaces are used to initially encode information and to maintain the same information in working memory^11,13–17^. But it is generally assumed that a given subspace uniquely codes for a specific type of information within a specific time frame. However, most studies use simple working memory tasks which vary only a single variable and analyses that focus on one task variable at a time.

Here, we use complex multi-factor tasks and multi-factor analyses to show that flexibility can even be seen in the subspaces themselves. Subspaces used for one purpose can be reactivated later for other purposes and can even be recycled to code for other types of information. This suggests that subspaces act more like generalized workspaces that may be optimized for different computations rather than being tied to representation of specific types of information.

## Results

### Object-Response association task and analysis

We reanalyzed data from a conditional visuomotor associative learning task that required integrating object information with an internal rule to generate a correct response (“Object task”)^18,19^. One of two object cues (’A’ or ‘B’) was displayed, followed by a blank delay and finally a “go” cue (Fig. 1A). In some blocks of trials, object A instructed a leftward saccade, and object B a rightward saccade. In alternating blocks, the response contingency rule was reversed, such that A instructed a rightward saccade, and B a leftward (Fig. 1B). The reversed task rule was relearned through trial and error after each unsignaled block switch.

**Figure 1.**
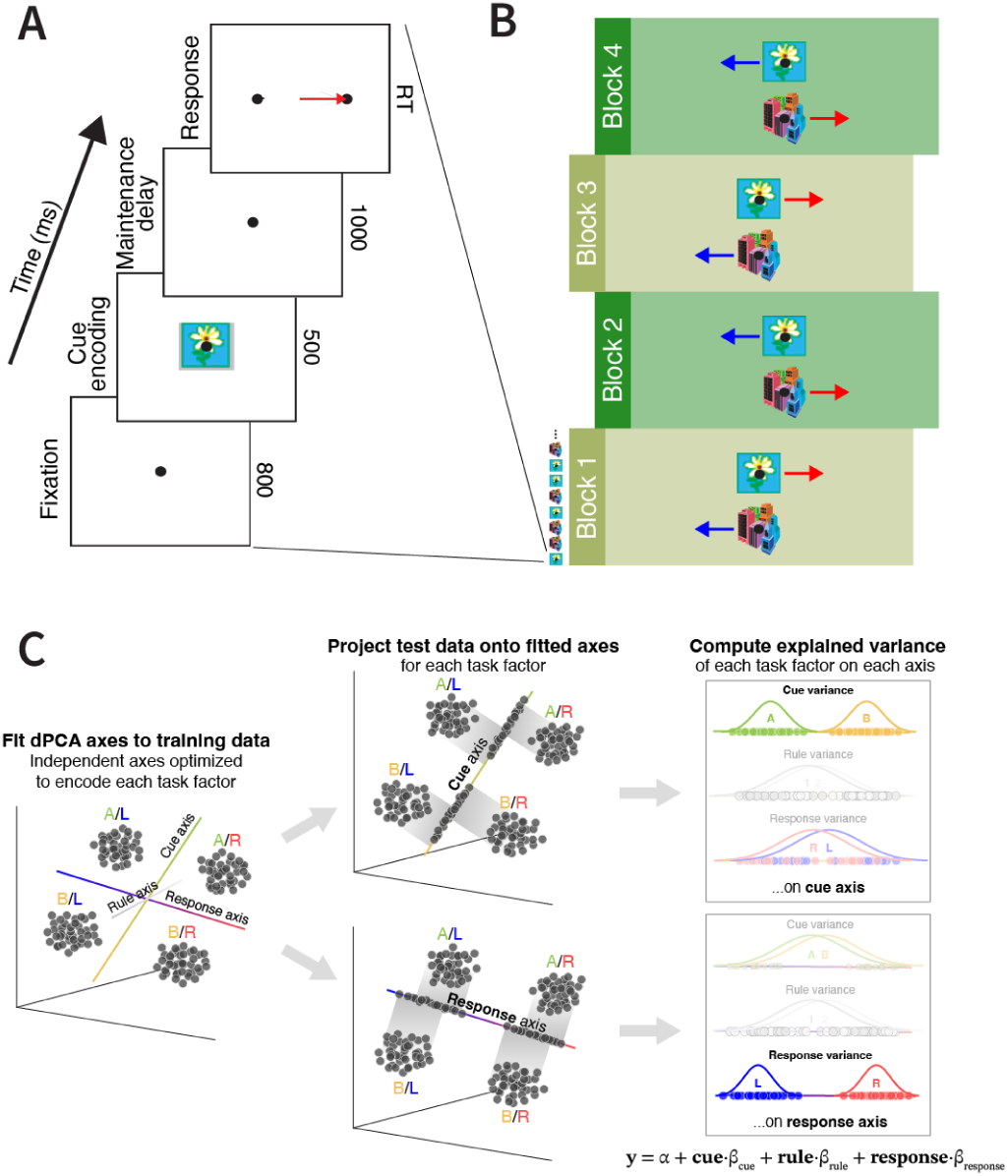
Object-Response association task. (A) Trial sequence. In each trial, one of two cue objects was shown. Each object instructed a saccade to one of two response targets, with a learned arbitrary mapping. (B) Task design. In alternating trial blocks, the object-response mapping was reversed, and subjects had to relearn the reversed mapping. (C) Data analysis strategy. Left: dPCA estimates axes (i.e. a subspace) within population activity space that maximize data variance for each task factor while minimizing variance for all others. Center: Independent test data is projected onto each of these axes (task rule axis not shown for conciseness). Right: A regression model quantifies how much test data variance along each axis can be explained by each task factor. By design, factors other than the one used to fit the dPCA axes (semi-transparent) should show little explained variance (i.e. the factors are “demixed”; confirmed in Fig. S1).

Previously we analyzed learning^18^ and reward coding^19^ in this task. Here, we focus on neural coding after each association was well learned in each rule block (final 15 trials of each condition per block). Examining post-learning data across both mapping rules resulted in data where the object cue and instructed response were orthogonal, so that we could disambiguate their neural codes.

Spiking activity was recorded from 632 neurons in the PFC of two non-human primates (NHPs). We developed an analysis pipeline that quantified information within population activity subspaces reflecting each task factor (Fig. 1C). We used demixed principal components analysis (dPCA)^20^ to identify axes in population activity space that maximize data variance for each task factor while minimizing variance for all others. This resulted in a subspace for each factor. We then projected cross-validated data onto these axes, and regressed the projected data onto all the task factors. This pipeline allowed us to determine how much information (quantified as the percent of projected data variance explained) about other factors “leaked” across subspaces. When this analysis was run independently at each time point, we found little leakage (Supplementary Fig. S1), indicating dPCA had successfully demixed the factors. Within their own subspaces, however, there was robust information about both the cue and the behavioral response (Supplementary Fig. S1).

### The PFC switched back and forth between subspaces

We examined whether the subspace PFC used to represent the same information was consistent over trial epochs or changed dynamically. We estimated dPCA axes at each time point, then quantified how much task information was conveyed in test data projected onto them from every other time point (Fig. 2A and Methods)^13,14,21^. This produced a cross-temporal “subspace generalization matrix” analogous to the familiar cross-temporal decoding matrix^13–17^. The vertical axis shows the time at which the subspace was estimated. The horizontal axis shows the time from which the data was projected into the subspace.

**Figure 2.**
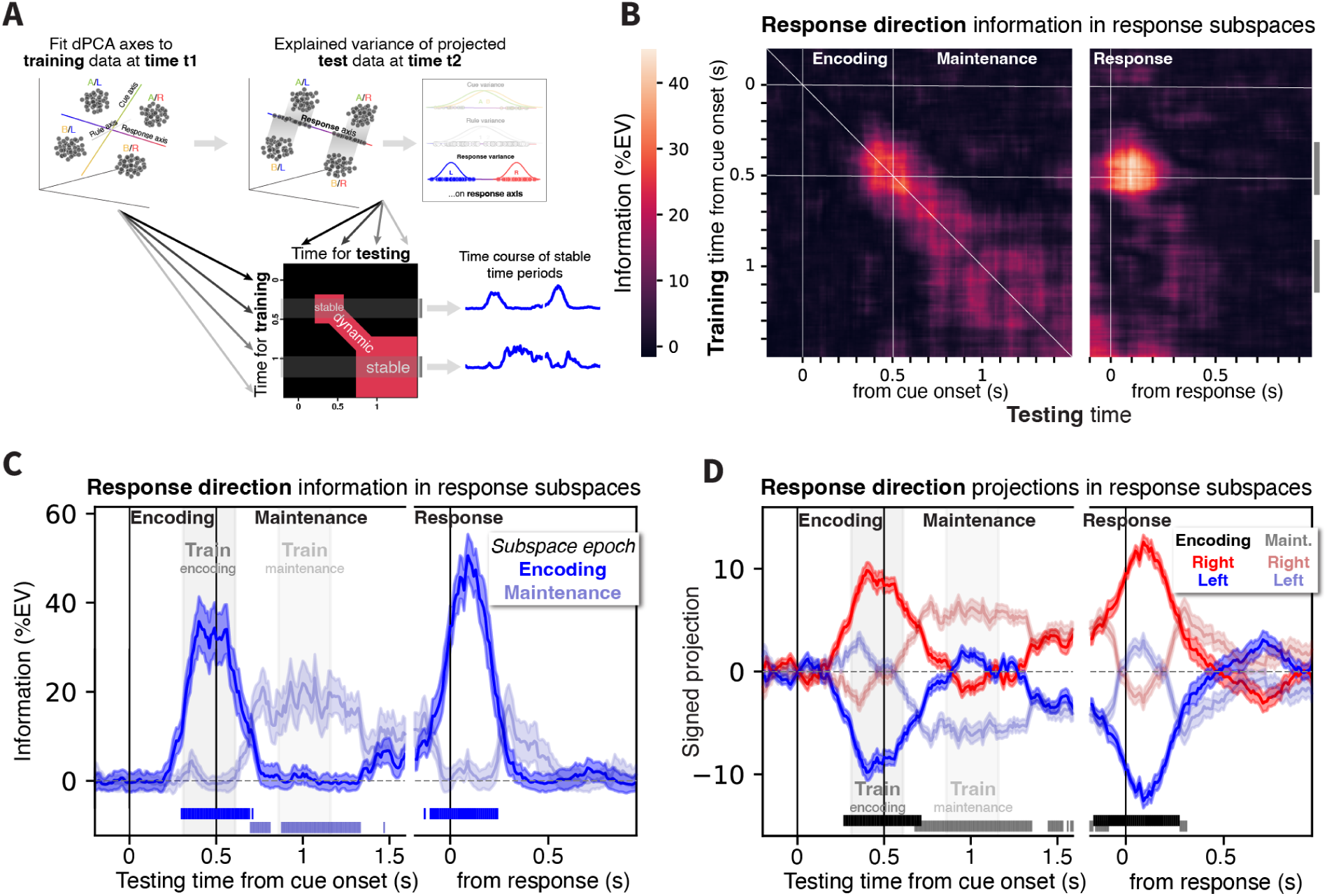
Reactivation of prefrontal subspaces. (A) We generalized our analysis strategy (Fig. 1C) to fit dPCA axes to training data from one time point, then measure information in projections of test data from a distinct time point. By training and testing at all pairs of time points, we generated a cross-temporal subspace generalization matrix, where each value reflects the percent of variance explained (%EV) at a given test time point (x-axis), when projected onto a dPCA axis fit at a distinct time point (y-axis). The matrix diagonal reflects data trained and tested at the same time (cf. Fig. S1). Off-diagonal points reflect the degree to which a subspace from one time point also conveys the same type of information at other times. (B) Cross-temporal subspace generalization matrix for PFC information about response direction in subject A (see Fig. S2 for results and significance for all task factors; similar results were found in subject P; Fig. S3). The PFC population dynamically shifts between locally-stable subspaces during the cue epoch (“encoding”; upper-left) and delay epoch (“maintenance”; lower-left). Unexpectedly, the encoding subspace is reactivated at the behavioral response (upper-right). (C) Summary of response direction information in data projected onto axes fit during encoding (darker curve and fill) and maintenance (lighter). (D) Mean signed projection onto encoding (darker) and maintenance (lighter) axes for rightward (red) and leftward (blue) response trials. Reactivation of the encoding subspace during the behavioral response has the same sign as its initial activation.

We found different subspaces for encoding and maintenance of information in working memory, consistent with previous studies using cross-temporal decoding^11,13–17^. Information about the behavioral response first appeared during the cue epoch (Fig. 2B). But this “encoding subspace” did not generalize well over the memory delay: data from the delay epoch no longer carried information about the behavioral response when projected into the encoding subspace (Fig. 2C, dark blue line). Instead, information appeared in a new subspace during the memory delay. This “maintenance subspace” carried behavioral response information that generalized over most of the delay epoch (Fig. 2C, light blue line). It did not generalize back to the cue epoch nor forward to the behavioral response epoch. Comparison of the angles of their dPC axes confirmed that these two subspaces were approximately orthogonal (Supplementary Fig. S4). These results indicate the same task factor—response direction—was encoded in two distinct subspaces: one during encoding and another when it was maintained in working memory. Near the end of the delay, response information in the maintenance subspace decreased, disappearing during the response epoch (Fig. 2C).

At the same time, however, information about the behavioral response *reappeared* in the initial encoding subspace (Fig. 2C). That is, spiking data from the response epoch carried strong information (explained variance) about the response direction when projected into the subspace estimated from the cue epoch. This effect began before the start of the response and continued after it. This indicates a strong overlap between the neural codes used to represent the behavioral response during the encoding and response epochs. This was confirmed by the small angle between their axes (Supplementary Fig. S4). These results suggest that, following the transition from the encoding to maintenance subspace, the encoding subspace was reactivated when it was time to read out the information to execute the behavioral response.

Furthermore, the behavioral response was represented with a consistent mapping during its initial appearance and subsequent reappearance in the encoding subspace. The previous results alone cannot capture this because they only relate to the *magnitude* of projections into a subspace. If the reappearance truly reflects a reactivation of a previous subspace, information should also be represented in the same way after reappearance. In other words, the *sign* of the projections should also be preserved. This was indeed the case. Figure 2D shows condition-mean activity for the right and left behavioral response trials over time, when projected into the encoding subspace (Fig. 2D, darker colors). When information reappeared in the encoding subspace after the delay, the projections were in the same direction as their initial appearance. This indicates consistent mapping of the two response directions. The same analysis is also shown for the maintenance subspace (Fig. 2D, lighter colors). These results suggest the reappearance of the encoding subspace reflects a reactivation of a previously utilized subspace in a later trial epoch.

In contrast, PFC population coding for the cue object showed only partial separation into encoding and maintenance subspaces, and no subspace reactivation (Supplementary Fig. S2C,D). This suggests that subspace reactivation is not a universal phenomenon, but may be restricted to coding of a behavioral response.

The results presented above came from one NHP. A second NHP showed effects with similar, albeit non-significant, trends (Supplementary Fig. S3). This could be due to unobserved differences in cognitive strategies by the NHPs. Thus, to confirm the subspace reactivation effect in another NHP, we turned to a different experiment that also involved cue-response associations but had another factor, spatial context. This also allowed us to test further hypotheses about subspace coding.

### Object/Location-Response association task

We analyzed data from a previously unpublished conditional visuomotor association task that required integration of a cue’s object identity *and its location* to generate the correct behavioral response (Fig. 3A,B). One of four possible object cue images (’A’, ‘B’, ‘C’, or ‘D’) was displayed at one of four possible locations (upper-right, upper-left, lower-left, or lower-right; Fig. 3A). Here, the associative mapping rule was constant over time, but reversed depending on the *location* of the object cue (Fig. 3B). When it appeared in the upper-left or lower-right locations, object A or B instructed a leftward saccade, while C or D instructed a rightward saccade. In the upper-right and lower-left locations, the mapping was reversed. Thus, this task required integration of object and spatial information (“what” and “where”) to generate a correct response. After the task was well-learned, spiking activity was recorded from 509 PFC neurons in an NHP subject.

**Figure 3.**
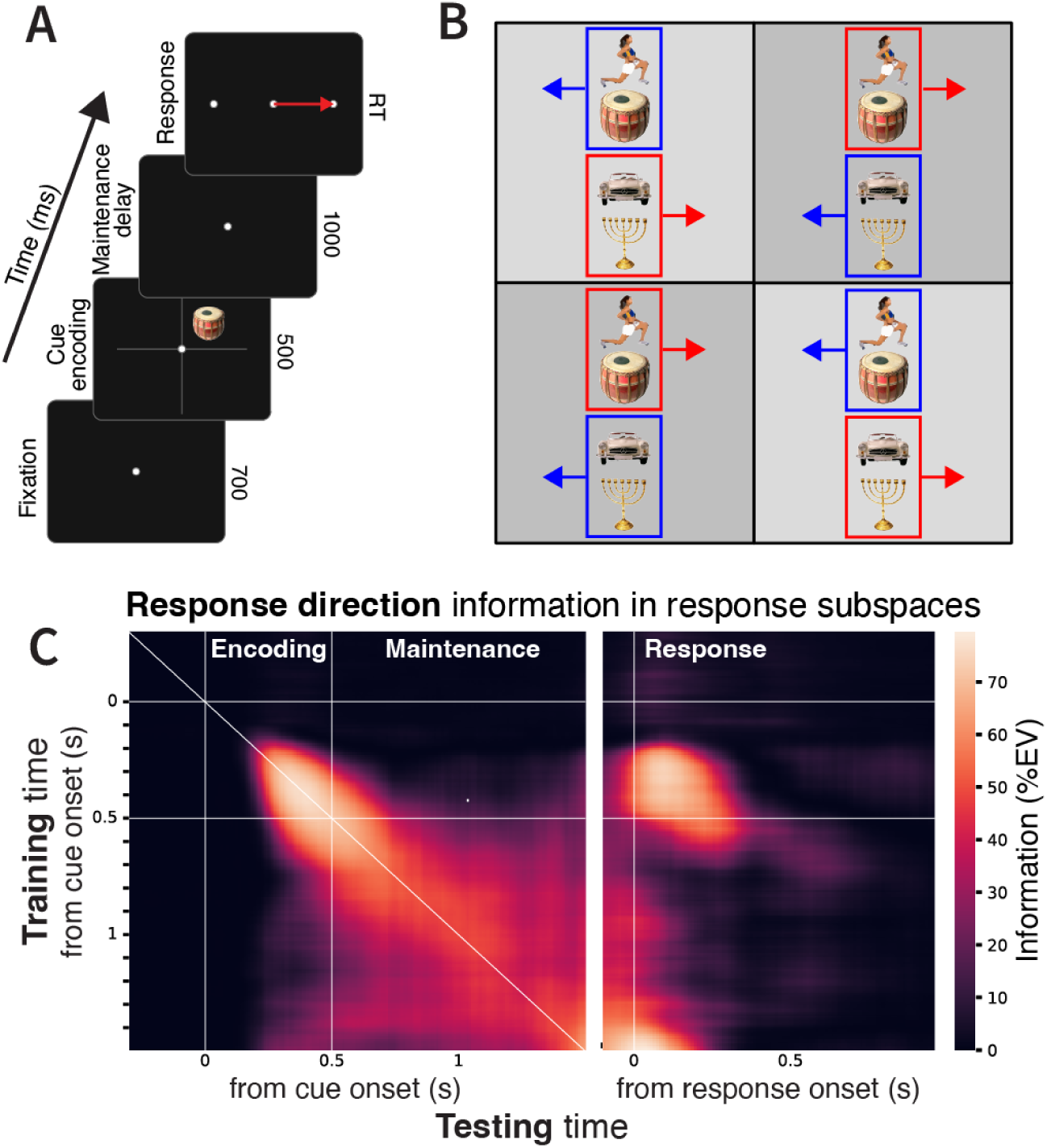
Subspace reactivation in an Object/Location-Response association task. (A) Trial sequence. Each trial, one of four cue objects was displayed in one of four locations. Following a blank delay, subjects made a saccadic response instructed by the specific combination of object and location. (B) Task design. Object-to-response mapping reversed depending on object location. In the upper-right and lower-left locations, two of the objects instructed rightward responses, while the other two objects instructed leftward responses. In the upper-left and lower-right locations, these same objects instructed the opposite response directions. (D) Cross-temporal generalization of PFC subspaces for response direction. PFC exhibits locally stable subspaces during the encoding and maintenance epochs, but the encoding subspace is reactivated during the behavioral response (upper-right).

### PFC subspace switching and reactivation in the Object/Location task

Subspace switching and reactivation was also observed in this task. Similar to the Object task, we observed an “encoding” subspace representing the behavioral response during the cue epoch. This was followed by a switch to a “maintenance” subspace that appeared after cue offset and persisted through the delay epoch (Fig. 3D). As in the previous task, these two subspaces were nearly orthogonal (Supplementary Fig. S6) and there was little generalization between them. That is, data from each time epoch projected into the other’s subspace carried near-zero information about the behavioral response (Fig. 3D).

As observed in the previous task, the encoding subspace—used initially during the cue epoch—was reactivated when the subject made a behavioral response (Fig. 3D, upper-right; Fig. 3E). This reactivation effect again began before the start of the response and continued after it, and again had the same mapping as in the initial activation (Fig. 3F). Thus, the results from two different tasks indicate that PFC can switch between subspaces and reactivate a previously utilized subspace to convey the same information.

### PFC subspaces for visuospatial location are recycled to encode response direction

The previous results showed that a PFC subspace can disappear and be reactivated in a later trial epoch to represent the *same* information. Here, we will show that a subspace encoding one type of information can also later be recycled to represent a *different* type of information. In the Object/Location task, the cue object’s identity must be integrated with its location. This means that both the behavioral output (response direction) and one aspect of the sensory input in this task could both be defined in visual space. We will show that the subspaces used initially to represent the cue location were subsequently used to represent the direction of the behavioral response during the response epoch.

Before examining cross-factor subspace generalization, we characterized the coding of cue location itself. Similar to results above for other task factors, we found that cue location information was again represented in independent subspaces during the encoding vs maintenance epochs, with no generalization between them (Fig. 4A, Supplementary Fig. S6). Neither subspace carried any information about cue location during the response period (Fig. 4A, right).

**Figure 4.**
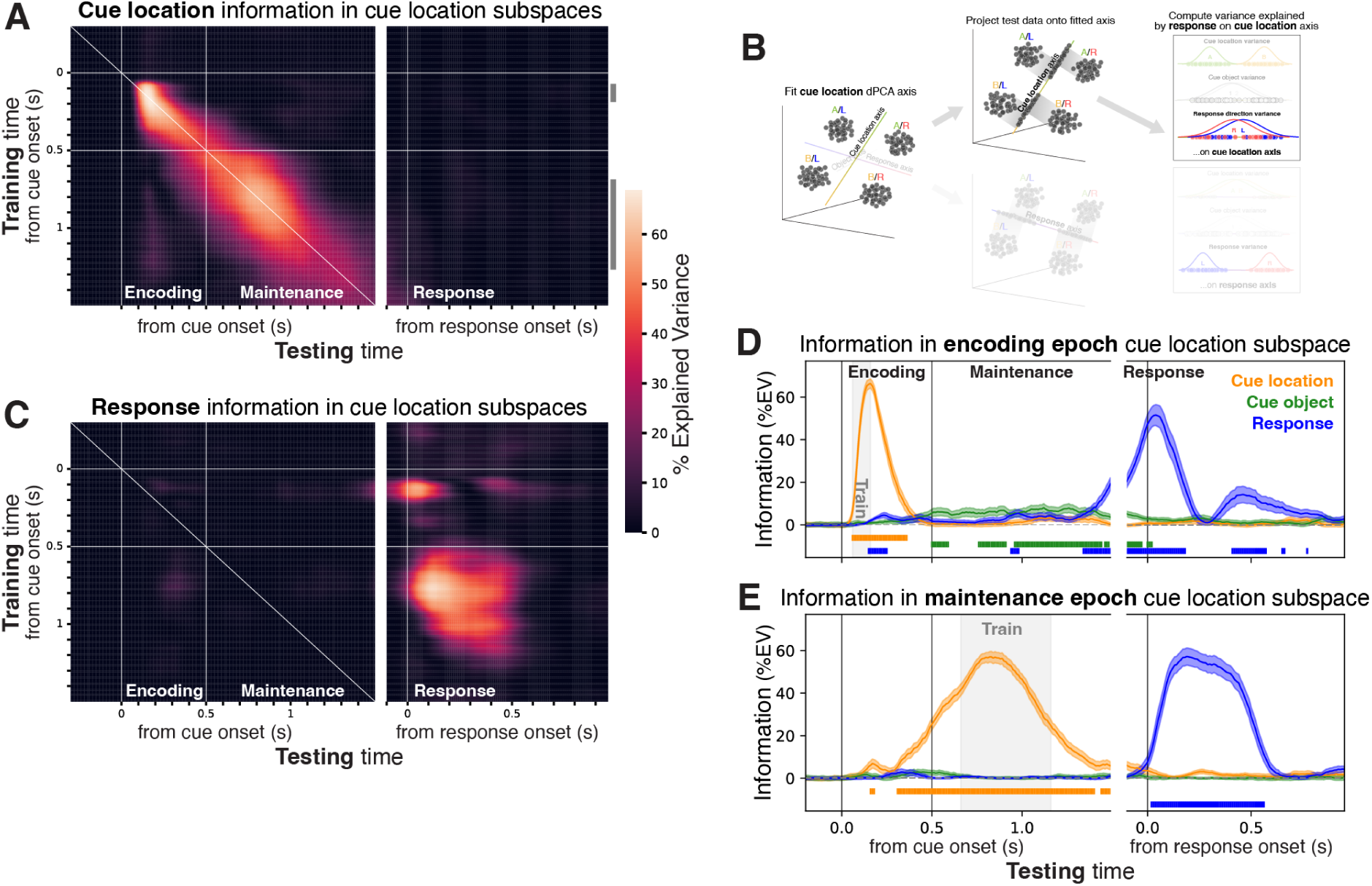
Prefrontal subspaces can be recycled to encode different types of information. (A) Cross-temporal generalization of PFC cue location subspaces. Color axis reflects variance explained by cue location at a given test time (x-axis) projected onto the cue location dPCA axis at a distinct training time (y-axis). Similar to our other results, there is a dynamic shift from locally stable subspaces during the encoding (upper-left) and maintenance epoch (lower-left). Neither subspace generalized to any times during the response epoch (right). (B) Cross-factor dPCA/regression analysis. We fit a dPCA axis to explain one task factor (cue location), then tested whether data projected onto it from other time points carried information about a distinct task factor (response direction). (C) Cross-factor generalization of cue location subspaces to encode behavioral response direction. Color axis reflects variance explained by response direction, for test data from a given time (x-axis) projected onto a cue location subspace trained at a distinct time (y-axis). Cue location axes contained no information about the behavioral response during the cue and delay epochs (left). But subspaces trained to reflect cue location earlier in the trial later carried information about the response during the response epoch (right; see Fig. S5 for significance and results for all pairs of task factors). (D–E) Summary of response direction information in data projected onto subspaces for cue location during the encoding (D) and maintenance (E) epochs.These results indicate prefrontal subspaces apparently dedicated to representing spatial sensory information can later be recycled to encode the behavioral response.

A key advantage of our dPCA/regression analysis is that we can also examine cross-factor generalization of subspaces. That is, we can train a subspace to reflect one factor and then measure the information within that subspace about *another factor* (Fig. 4B). We used this to see whether projections onto the cue location axes at each time point also carried information about behavioral response direction. This produced a *cross-factor* subspace generalization matrix (Fig. 4C). The vertical axis reflects the time at which a subspace was trained to represent *cue location*. The horizontal axis reflects the time from which data was projected into a cue location subspace to estimate information about *response direction* (Fig. 4C). Note that these two factors were orthogonal in the task design and their representations had no overlap at the times when the cue location subspaces were estimated (diagonal in Fig. 4C), reflecting successful demixing by the dPCA.

Nevertheless, the cue location “encoding” subspace—which, again, was trained to reflect only cue location during the cue epoch—later conveyed strong information about behavioral response direction during the response epoch (Fig. 4, upper-right; Fig. 4D, blue line). This effect began near the end of the delay epoch and continued into the response epoch. It also showed a smaller “rebound” of reactivation after the behavioral response (Fig. 4D, at ∼0.5 s). These results indicate that a PFC subspace originally used for representing visuospatial location can later be recycled to encode behavioral response direction.

Further, the cue location “maintenance” subspace—again, trained to reflect only cue location during the delay epoch—was also recycled to convey the behavioral response later, after the response (Fig. 4C, lower-right; Fig. 4E, blue line). This occurred after the first peak of recycling for the cue location “encoding” subspace (Fig. 4E). This suggests that subspace recycling during the response epoch recapitulates the shift from encoding to maintenance subspaces. We note that a similar, but nonsignificant, trend was observed for simple reactivation of the “maintenance” subspace for the behavioral response (Fig. 2C,3C). These results provide evidence that neural subspaces—and entire dynamic sequences of subspaces—can be recycled for different purposes.

Note that these results do *not* simply mean that cue location and response direction were collapsed into a single unitary variable in the PFC representation. Neither the encoding or maintenance subspaces for cue location contained any information about the behavioral response during the cue or delay epochs (Fig. 4C, left). Instead, during these epochs, response direction was encoded in its own distinct set of subspaces (Fig. 3C). Further, during the response epoch, when the cue location subspaces were recycled to convey information about the behavioral response, they no longer carried information about the cue location (Fig. 4A, right). Thus, cue location and response direction information used different subspaces during the encoding and maintenance epochs. But during the response epoch, the behavioral response “co-opted” the subspaces previously used to encode cue location.

### PFC subspaces used congruent spatial maps to represent cue location and response direction

In the previous section, we showed that cue location and response direction used the same subspaces, albeit at different times. Here, we asked whether these two usages of the same subspaces had congruent visuospatial mappings. That is, did “left” vs “right” in cue location subspaces correspond to leftward vs rightward responses when they were recycled for the behavioral response?

To analyze this, we wanted to estimate a transformation of the 2D cue location subspace where the subspace dimensions specifically corresponded to standardized left-right and up-down axes (Fig. 5A). We fit two linear classifiers to optimally categorize object locations along the two diagonals (lower-left vs. upper-right and upper-left vs. lower-right; Fig. 5A(i)). This resulted in axes that were nearly orthogonal, but with a bias toward an expanded representation of the positive diagonal (Fig. 5A(ii)). We thus orthogonalized and scaled these axes to unit length. To produce a standard Cartesian coordinate system that reflects the spatial mapping of object location in prefrontal activity, we rotated the orthogonalized axes by –45° (Fig. 5A(ii)). Finally, we projected population activity for left vs right response directions onto these axes to determine how the behavioral response maps onto the prefrontal cue location space (Fig. 5A(iv); see Methods for details).

**Figure 5.**
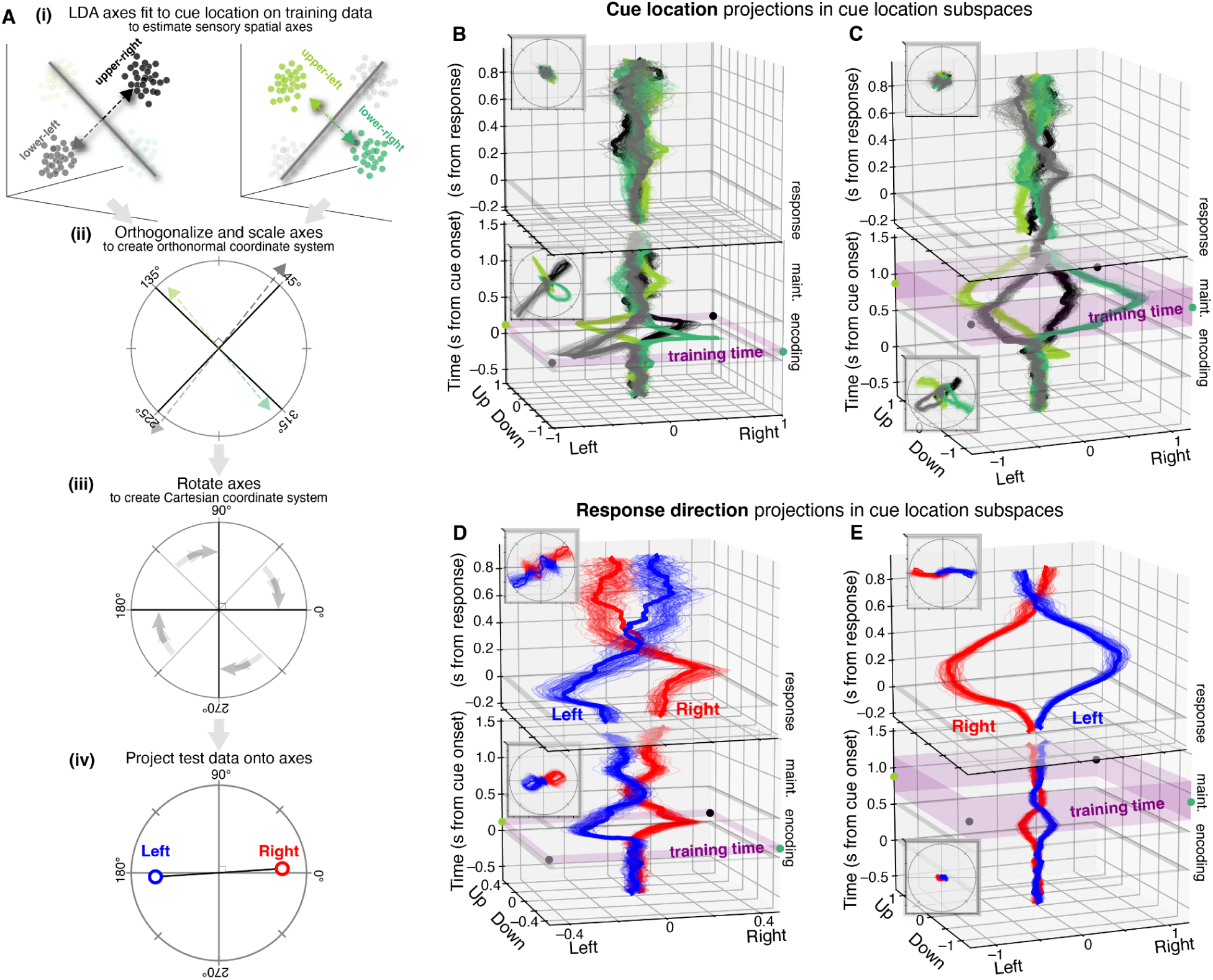
Prefrontal subspace recycling follows the mapping of the recycled space. (A) Analysis strategy to map response direction onto neural space for 2D object location. Two linear discriminant axes were fit to categorize object locations along the positive and negative diagonals in training data. The resulting axes were orthogonalized, scaled, and rotated to create a standardized Cartesian coordinate system. Cross-validated left and right response trials were projected onto these axes across time within trials to estimate their location in object location space. (B,C) Cross-validated projections across time of condition means for four object locations onto 2D object location space trained during encoding (B) and maintenance (C) epochs (purple fill: fitting time window), for 100 bootstrap resamples (thin lines; thick lines: mean across bootstraps; insets: top views). Object cue locations project to expected locations during the training epochs, but have little projection during the response epoch, confirming method validity. (D) Projections of left (blue) and right (red) response direction trial means onto cue location space trained during encoding epoch. Responses lie approximately in their appropriate locations in cue location space during the peri-response epoch, but reverse to approximately opposing locations during the post-response epoch. (E) Projections of response direction onto maintenance-epoch cue location space. Post-response projection is reversed. Results show that response direction recycles cue location subspace in a way roughly consistent (or anti-consistent) with the organization of its initial 2D mapping.

To first validate our method, we projected the (cross-validated) cue locations onto these axes (Fig. 5B,C). Projections across time (vertical axis) into this space are shown for the left-right and up-down axes (insets show “birds-eye” views of projections with only the spatial axes). As expected, cue locations projected to their appropriate places at the four quadrants (Fig. 5B,C, bottom), though with some remaining bias toward an expanded representation at the lower-left location. As in previous results (Fig. 4), there was little evidence of any projection of cue location onto these axes during the response epoch (Fig. 5B,C, top).

We next examined the projection of response direction onto the cue location subspace, our main question of interest. We found that behavioral response direction information during subspace “recycling” roughly matched the original mappings in cue location subspace. That is, left and right responses projected roughly to where “left” and “right” would be, respectively, in the cue location subspace (Fig. 5D). This suggests that response direction not only recycles cue location subspace, but does so in a way that broadly preserves its spatial mapping. The modest counter-clockwise rotation away from horizontal may be related to the remaining over-representation of the lower-left location seen during coding of cue location (Fig. 5B). Interestingly, for the weaker secondary post-response recycling of the encoding-epoch subspace, the mapping is almost exactly reversed—left and right responses projected roughly to “right” and “left” in the cue location space. This same approximate direction reversal is observed when response directions are projected onto the maintenance-epoch cue location space (Fig. 5E). These post-response direction reversals might be related to termination of processes that lead to the response saccade or to planning/execution of the return saccade in the opposite direction to shift gaze back to the center for the next trial.

### Cue object subspaces are also recycled to encode behavioral response direction

Thus far, we have shown evidence for recycling of a subspace initially dedicated to cue location to later encode response direction, another spatial variable. However, both tasks also included a *non-spatial* variable, the cue object identity. We wondered if the recycling of different subspaces seen above was due to their shared use of the spatial dimension or whether non-spatial subspaces could also be recycled to reflect the response direction. This was not the case for the Object task presented at the start of the Results. Behavioral response direction showed no significant projection onto subspaces for the cue object (Supplementary Fig. S2,3).

However, a different picture emerged from the Object/Location task. There were again distinct subspaces for cue object identity during the encoding and maintenance epochs (Fig. 6A, left; Supplementary Fig. S6). The short-lived encoding subspace for the cue object showed little evidence of carrying information about other task factors at any time point (Fig. 6B–D). In contrast, the maintenance subspace for the cue object showed a complex triphasic pattern of activation and reactivation (Fig. 6B,C,E, Supplementary Fig. S5). It was only trained to encode the cue object during the maintenance epoch (Fig. 6E, gray rectangle). However, this subspace represented the cue location during the encoding epoch (Fig. 6E, orange) and then response direction during the response epoch (Fig. 6E, blue). As with previous results, in each of these epochs, information about only a single task variable predominated in this subspace, and the others essentially disappeared. These results show that PFC can recycle non-spatial subspaces (cue object) to encode spatial information (cue location and response direction).

**Figure 6.**
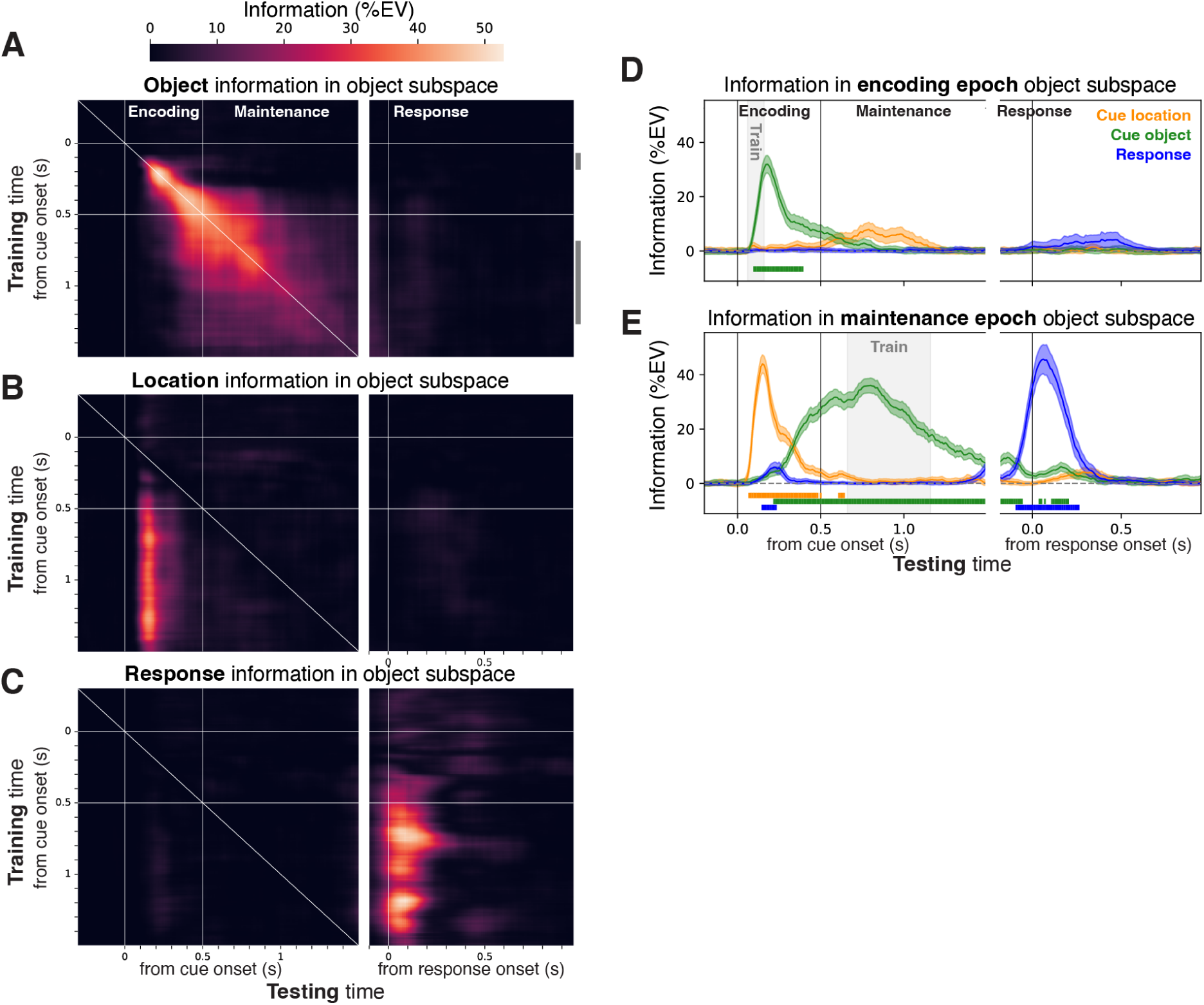
Prefrontal subspace for cue object identity is recycled to represent cue location and response direction. (A) Cross-temporal generalization of PFC cue object subspaces. Color axis reflects variance explained by object identity at a given test time (x-axis) projected onto the object dPCA axis at a distinct training time (y-axis). Again, there is a briefly stable subspace during the encoding epoch (upper-left) and a more temporally-extended epoch of stability in the delay (lower-left). Neither subspace generalized to any times during the response epoch (right; see Fig. S5 for significance). (B) Cross-factor generalization of cue object subspaces to encode cue location across subspace training (y-axis) and testing (x-axis) times. Heatmap color reflects variance explained by cue location, for test data projected onto the cue object subspace. Object subspace during the maintenance epoch conveyed information about cue location during the early encoding epoch. (C) Cross-factor generalization of cue object subspaces to encode response direction. Color reflects variance explained by response direction, for test data (x-axis) projected onto the cue object subspace (y-axis). Object subspace during the maintenance epoch also conveyed information about response direction during the peri-response epoch. (C,D) Summary of time courses of explained variance for all task factors, for the stable object subspaces during the encoding epoch (C) and maintenance epoch (D). While the encoding-epoch space showed little evidence of recycling, the delay-epoch exhibited a complex temporal pattern of recycling. It initially conveys information about cue location (orange), then cue object (which it is optimized for; green), then finally it conveys information about response direction (blue). These results indicate that even non-spatial PFC subspaces can be recycled to carry other types of information, including multiple different types of information in complex temporal sequences.

## Discussion

Our results suggest that subspaces can be flexibly used for different purposes. Representation of the behavioral response moved from one subspace when it was initially encoded to an orthogonal subspace during memory maintenance. The original subspace was then reactivated during the behavioral response itself (Fig. 7A). When both a cue object’s identity and its location instructed a behavioral response, we found that subspaces initially representing sensory information were recycled later to represent the behavioral response (Fig. 7B). Recycling of a subspace preserved its original spatial organization. In other words, “cue right vs left” in the original sensory subspace was also “response right vs left” when it was recycled for the behavioral response. These results suggest that PFC subspaces are highly flexible, and activity within them can “mean” many different things at different times.

**Figure 7.**
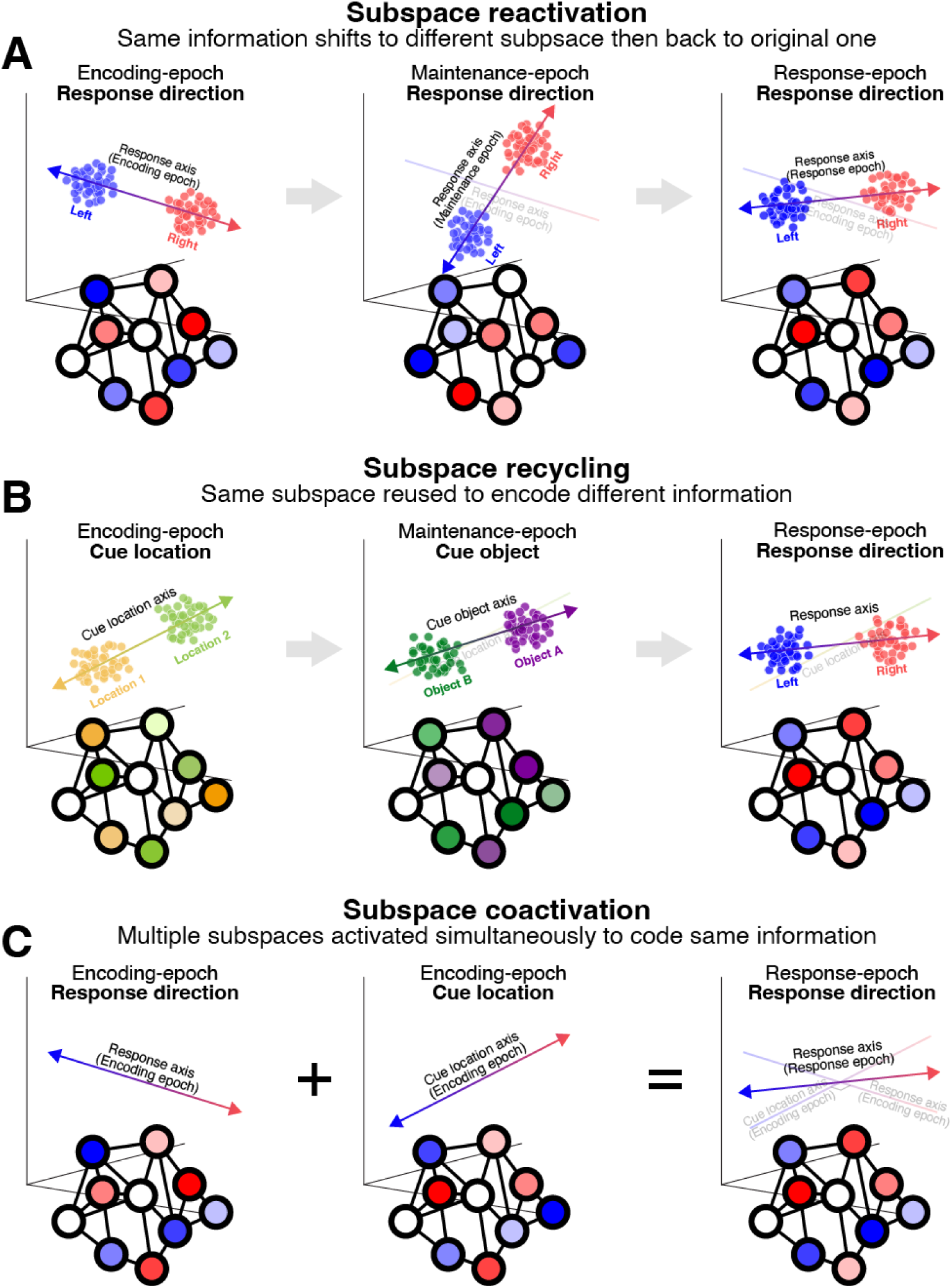
Summary of key results. (A) Subspace reactivation. The subspace PFC used to code for the behavioral response direction changed from the encoding to maintenance epoch. But the encoding-epoch subspace was reactivated to represent response direction during the response itself. These results suggest information can be rotated out of a subspace, then later back into it again. (B) Subspace recycling. The same subspace that codes for the cue location (during the encoding epoch) later codes for the cue object identity (maintenance epoch), then the response direction (response epoch). These results suggest PFC can recycle the same subspaces to reflect different types of information. (C) Subspace coactivation. Results are consistent with coactivation of both the encoding-epoch response direction subspace (A, left) and the encoding-epoch cue location subspace (B, left) to code for response direction during the response epoch. This results in a population activity pattern that is a mixture of both components, and a population axis that is in between the corresponding component axes.

We also find evidence that multiple subspaces that initially seem distinct can come to “mean” the same thing. Subspaces coding for the cue location and behavioral response direction during the encoding (cue) epoch were approximately orthogonal. But when response-epoch activity was projected onto these subspaces, both conveyed information discriminating right vs left responses. When we trained a subspace to specifically reflect response direction during the response epoch, it was partially correlated with both of these encoding-epoch subspaces. These results are consistent with a situation where multiple orthogonal subspaces are activated simultaneously (Fig. 7C). If neural population patterns corresponding to two orthogonal axes in activity space are coactivated, the result will be a weighted vector sum of them. That is, a third axis in between the two coactivated ones, which will have strong projections onto both of them. Thus, our results are consistent with the possibility that PFC activity during the response epoch reflects coactivation of both of these encoding-epoch subspaces to code for the behavioral response.

Our results showing flexible use of different subspaces expand on previous studies showing that neural coding of working memory is dynamic over short timescales^11,13–17^. These studies used decoding and related methods to show that population activity patterns reflecting the same information change over the course of a trial. However, these studies only demonstrated that such activity patterns change in one direction across time (e.g. encoding→maintenance), and convey a single type of information. By contrast, our subspace analysis indicates that activity patterns are not simply used once and discarded. Subspaces can be reactivated later for a different purpose (e.g. encoding vs. behavioral response) and can even be recycled to represent different types of information. Analogously, the leading models of these dynamics are recurrent neural networks with asymmetric weights that add effectively feedforward interactions to within-PFC recurrent connections^17^. Our results suggest these models should be updated to account for the reactivation of previous subspaces, and extended to account for subspace recycling.

Adaptive coding theories of prefrontal function emphasize the ability of PFC to flexibly convey task-relevant information to guide input/output mappings from sensory to motor cortex^2,4^. A key feature of this flexibility is that individual neurons multiplex various types of information, a phenomenon termed “mixed selectivity”^3,5,21,22^. Multiplexing entails an ambiguity in the sense that a single neuron’s spike rate could “mean” many different things. Prior work suggests that such ambiguity is resolved through population coding, where distinct patterns of population activity correspond uniquely to the information they code for^3,6–8^. Our results extend this framework by showing that information can even be multiplexed at the level of subspaces. We found that similar patterns of population activity (defining a subspace) can represent different information at different times. This suggests that unambiguous representation of specific information may only emerge at the level of dynamic activity streams spanning entire brain systems.

In addition, these results suggest that prefrontal subspaces may be more like workspaces where different types of information can be “written”. Rather than being tied to the information they represent, subspaces might be better characterized by the computations they enable. For example, the subspaces activated during both encoding and the behavioral response could reflect neural connectivity and activity patterns optimized for loading information into and reading it out of working memory^17^, or for online manipulation of information. In contrast, the subspaces activated during maintenance might be optimized for short-term storage of information. Rotating information into an orthogonal subspace might also gate its transmission to downstream areas, analogous to how motor cortex has been suggested to prevent movement^23,24^. If this latter possibility is true, it raises the question of why motor output was not triggered during the initial activation of the subspaces that were used during both encoding and the behavioral response. Perhaps the apparent coactivation of multiple subspaces we observed during the behavioral response (Fig. 7C) facilitates connection of working memory read-out to the motor system.

Our results indicate subspace *information content* seems to be quite flexible. On the other hand, a previous series of studies has shown that subspace *structure* appears to be relatively unmalleable. It is virtually impossible to learn to alter the population activity patterns that define subspaces themselves^25–27^. Taken together, these findings suggest that PFC, and possibly other cortical areas, contain only a limited set of subspaces that are reused as needed to encode different types of information. This is broadly consistent with the idea of a “global workspace” in working memory^28^. This theory suggests a single cognitive workspace that encodes whatever information is currently being used for cognition. Our results might be seen as an extension of this idea, where a limited set of workspaces are used to encode different types of information.

## Methods

### Non-human primate (NHP) subjects

Three rhesus monkeys (*Macaca mulatta*) were used in the studies reported here. For the Object task, two NHPs were used: NHP A was female, aged 6 years, 5 kg at the time of the study and NHP P was male, aged 5 years, 7 kg. For the Object/Location task, a single male NHP Z, aged 6 years, 10 kg was used. NHPs were pair-housed on 12 hr day/night cycles and maintained in a temperature-controlled environment (80° F). The NHPs were not involved in previous procedures. All procedures followed the guidelines of the Massachusetts Institute of Technology Committee on Animal Care and the National Institutes of Health.

### Task design

#### Object task

This task required integration of object identity and task rule information to generate a correct behavioral response (Fig. 1B). In each trial, NHP subjects maintained central gaze while one of two object cues (’A’ or ‘B’) was displayed at the fovea for 500 ms, followed by a blank delay epoch (1000 ms). Two response targets were then shown 6° to the left and right of fixation, and subjects were required to saccade directly to the correct target to obtain liquid reward. Object stimuli were arbitrarily mapped to response direction, with a mapping rule that was periodically reversed. In some blocks of trials, the task rule instructed a leftward saccade for object A and rightward for object B. In other blocks of trials, the response contingency was reversed—object A instructed right, and object B instructed left.

NHP subjects learned the initial task rule through trial-and-error reinforcement learning. Once they reached a behavioral criterion (90% correct over the previous ten trials for each cue), the response contingency was reversed with no explicit signal, and subjects had to relearn the new task rule again through trial-and-error. Each recording session consisted of 3–8 reversals (4–9 trial blocks). Object stimuli were complex multi-colored images from an image library (Corel Gallery photos). For each recording session, two never-before-seen images were selected at random.

#### Object/Location task

This task had a similar overall structure—subjects integrated multiple streams of information to compute arbitrarily associated behavioral responses. Here, rather than integrating sensory information with an internally-maintained task rule, subjects integrated two distinct types of sensory information—object and location.

In each task trial, the NHP subject maintained central fixation while one of four possible object cue images (‘A’, ‘B’, ‘C’, or ‘D’) was displayed for 500 ms in one of four possible locations (2° to the upper-right, upper-left, lower-left, or lower-right). To aid perception of the categorical object location, a cross stimulus was displayed simultaneously with it. Following a blank 1000 ms delay, two response targets were then shown 8.5° to the left and right of fixation, and subjects were required to saccade directly to the correct target to obtain liquid reward. Cue object was arbitrarily mapped to response direction, with a mapping rule that reversed depending on the object location. In the upper-left and lower-right locations, objects A and B instructed a leftward saccade and objects C and D instructed a rightward saccade. In the upper-right and lower-left locations, the opposite mapping was required—objects A and B instructed rightward saccade and objects C and D a leftward saccade. In this task, the same four well-learned objects were used across all sessions, and the mapping for a given object/location conjunction was never changed.

### Electrophysiological data acquisition

For both datasets, neural activity was recorded from ventrolateral and dorsolateral prefrontal cortex with dura-puncturing tungsten single electrodes (FHC Instruments). Electrodes were lowered acutely into the cortex for each daily session using custom-made, independently adjustable microdrives. The raw neural data was high-pass filtered and amplitude thresholds were set manually to extract neuronal spiking activity. For the Object task dataset, data was recorded using the Plexon MAP system, and spikes were manually sorted offline into isolated single neurons using amplitude and waveform features (Plexon Offline Sorter). For the Object/Location task dataset, data was recorded using the Blackrock Cerebus system, and spikes were not sorted into single neurons. Instead, all threshold crossings on each electrode were analyzed as multi-unit spiking activity. Eye position was monitored using infrared eye tracking systems (Object task: IScan,100 Hz; Object/Location task: Eyelink 1000, 1000 Hz).

### Data analysis

#### Data selection

##### Object task dataset

To ensure sufficient data for analysis, only sessions with a minimum of four completed blocks were included. An additional 11 neurons were rejected due to excessive artifacts from the solenoid controlling juice rewards. Only the last 15 correct trials from each completed trial block were included in the analyses. This was done to avoid including trials where the associative reversal learning was still occurring and focus analysis on trials where the association is well-learned. Otherwise, all neurons and all correct trials were included. These criteria resulted in a population of 318 neurons across 26 sessions in subject A, and 314 neurons across 22 sessions in subject P.

##### Object/Location task dataset

One recording session was removed due to a substantial response bias towards the right (351 vs 586). Otherwise, all units and all correct trials were included.

#### Preprocessing

For the Object task, spike rates were computed in 100 ms sliding windows, with a 10 ms step between them. Because temporal drifts in spike rates could be misinterpreted as task rule (trial block) effects in the Object task dataset, we estimated and removed them. Spike rates were smoothed with a broad across-trial Gaussian kernel (SD = 60 trials) to estimate slow trends in rate. This estimate was then subtracted from each individual trial, and the resulting drift-corrected rates were used for all further analyses. For the Object/Location task, spike rates were binned at 50 ms with a step size of 10 ms. The finer-grained temporal resolution was feasible due to higher signal-to-noise in this dataset, likely due to it not involving real-time learning.

For both datasets, data was collected from most neurons asynchronously. Thus, for both we used a pseudopopulation analysis strategy^29^. To generate each pseudopopulation, a random subset of trials from each task condition was selected for each neuron in the population (Object task: 30 trials for each of 4 conditions; Object/Location task: 33 trials for each of 16 conditions). Rates from the resulting pseudotrials were used for all subsequent analyses. To ensure results weren’t dictated by any specific random sample, and to generate bootstrap-like resamples for statistical analysis, pseudopopulation generation was randomly repeated 100 times.

#### Cross-validation

For all analyses, statistics of interest were estimated using 5-fold cross-validation. Selected trials within a pseudopopulation were split into five subsets. For each subset, a model (e.g. dPCA axes) was fit on 80% of trials. Then the statistic of interest was computed on the remaining 20% of trials in the test subset. This was repeated five times, with each of the five subsets taking a turn as the test subset. This same strategy was used for cross-temporal analysis except that, for each cross-validation split, the 80% of trials used for training came from one time point and the 20% used for testing came from a distinct time point.

#### dPCA-regression analysis

Demixed principal components analysis (dPCA)^20^ is a dimensionality reduction method that, like principal components analysis (PCA), finds axes in multivariate data (here, neural activity across a pseudopopulation) along which variance is maximized. But, like linear discriminant analysis (LDA), it is a supervised method and finds axes that maximize variance for a specific factor in a linear model characterizing the data, and minimize variance for all other factors and interaction terms. For the Object task, we estimated dPCA axes for object, task rule, and their interaction (corresponding to the response direction). For the Object/Location task, we estimated dPCA axes for cue object identity, cue location, and their interaction (corresponding to the response direction). Unlike many applications of dPCA, we did not include time as a dPCA factor. Instead, dPCA was run independently at each time point, so we could directly examine the temporal evolution of subspace coding without any assumptions about its correlation structure across time. dPCA axis estimation was performed on across-trial condition means computed from the training data subset.

To measure how much information a given dPCA axis conveyed about each task factor, we projected cross-validated single-trial data from the test subset onto the axis. The resulting 1D projection was then regressed onto a linear model containing all task variables. For the Object task, these included the cue object, the task mapping rule, and the response direction. For the Object/Location task, these included the cue object, cue location, and response direction. This was used to compute the percent of variance in the projected data explained by each task factor (both the factor used to estimate the dPCA axis and all others). For analyses of the sign of data projections (Fig. 2D), instead of the regression analysis, we simply computing the across-trial condition means of the projected test data.

This analysis is broadly similar to previous analysis methods, but has some key advantages. Like decoding analysis, it quantifies information conveyed by an entire neural population. But unlike decoding, our regression analysis considers all task factors together, properly dealing with correlations between factors, and not attributing variance to factors other than the decoded one to “noise” (see also Ref. 30). An advantage over previous methods quantifying multivariate population information^30^ is that explained variance is a familiar, well-normalized measure (ranging from 0–100). Finally, using dPCA instead of an unsupervised dimensionality reduction method (like PCA) has the advantage of discovering axes that are uniquely related to specific task factors and that minimize variance of all other factors. This property provides a crucial control to our results showing recycling of subspaces between different task factors (Fig. 4,6).

#### Subspace 2D mapping analysis

To quantify where the projections of left and right response directions fell in the cue location space (Fig. 5), we required a different analysis strategy (Fig. 5A). The previous analysis pipeline discovered a subspace whose axes captured maximal overall variance reflecting all four cue locations. Here, we desired a space whose axes specifically corresponded to the horizontal and vertical visual meridians.

We fit two linear discriminant axes (Fig. 5A, top): (1) classifying locations along the positive diagonal (*x*_+_; upper-right vs. lower-left), and (2) classifying locations along the negative diagonal (*x*_–_; upper-left vs. lower-right). Though there were no constraints enforcing orthogonality between these axes, they naturally were nearly orthogonal. To make them completely orthogonal, and thus a proper basis for projection, we used Löwdin symmetric orthogonalization^31^ (Fig. 5A, middle). This method orthogonalizes two or more axes without giving priority to any of them, as iterative methods like Gram–Schmidt do. The orthogonalized basis is given by *B*_*ortho*_ = *UV*^*T*^, where *USV*^*T*^ is the singular value decomposition of the original basis *B* = [*x*_+_, *x*_−_]. To equate the length of the axes, and reduce spatial biases in the population data, we scaled both orthogonalized axes to unit length. Finally, these orthonormal axes were rotated by –45° to generate a Cartesian coordinate system for visual space. Trials with left and right saccadic responses were projected onto the resulting axes and trial-averaged (Fig. 5A, bottom).

#### Statistical tests

All statistics used parametric bootstrap methods^32^. For all analyses, 100 pseudopopulations were generated by resampling of random trial subsets. Pseudopopulations were treated as bootstrap resamples, and each was used to estimate a statistic of interest (e.g. explained variance of a given task factor when data was projected onto a given dPCA axis). The mean and standard deviation across bootstraps were used as estimates of the mean and standard error of the statistic, and input into a *z*-test to compute *p* values. Effects of running multiple tests on the same data were corrected using the false discovery rate (FDR) method across all time points and task factors for each dataset.

#### Software

Data preprocessing was performed in MATLAB R2019b (The Mathworks, Inc, Natick, MA). All other analysis was performed in Python 3.10, using the scikit-learn, statsmodels, and spynal^33^ libraries, as well as custom code.

## Acknowledgements

We thank Morteza Moazami for collecting the Object/Location task dataset. We thank Adam Eisen and the Miller Lab for helpful comments and discussions on this work. This work was supported by Office of Naval Research N00014-22-1-2453, the Freedom Together Foundation, and The Picower Institute for Learning and Memory.

## Author contributions

A.P. and E.K.M. designed the experiments. A.P collected the data. S.L.B. curated the data. P.T.W. and S.L.B. designed and carried out the data analysis. P.T.W., S.L.B., and E.K.M. wrote the manuscript, and E.K.M. supervised the study.

## Resource availability

Requests for further information and resources should be directed to and will be fulfilled by the lead contact, Earl K. Miller.

## Supplementary Information

### Supplementary Results

**Figure S1.**
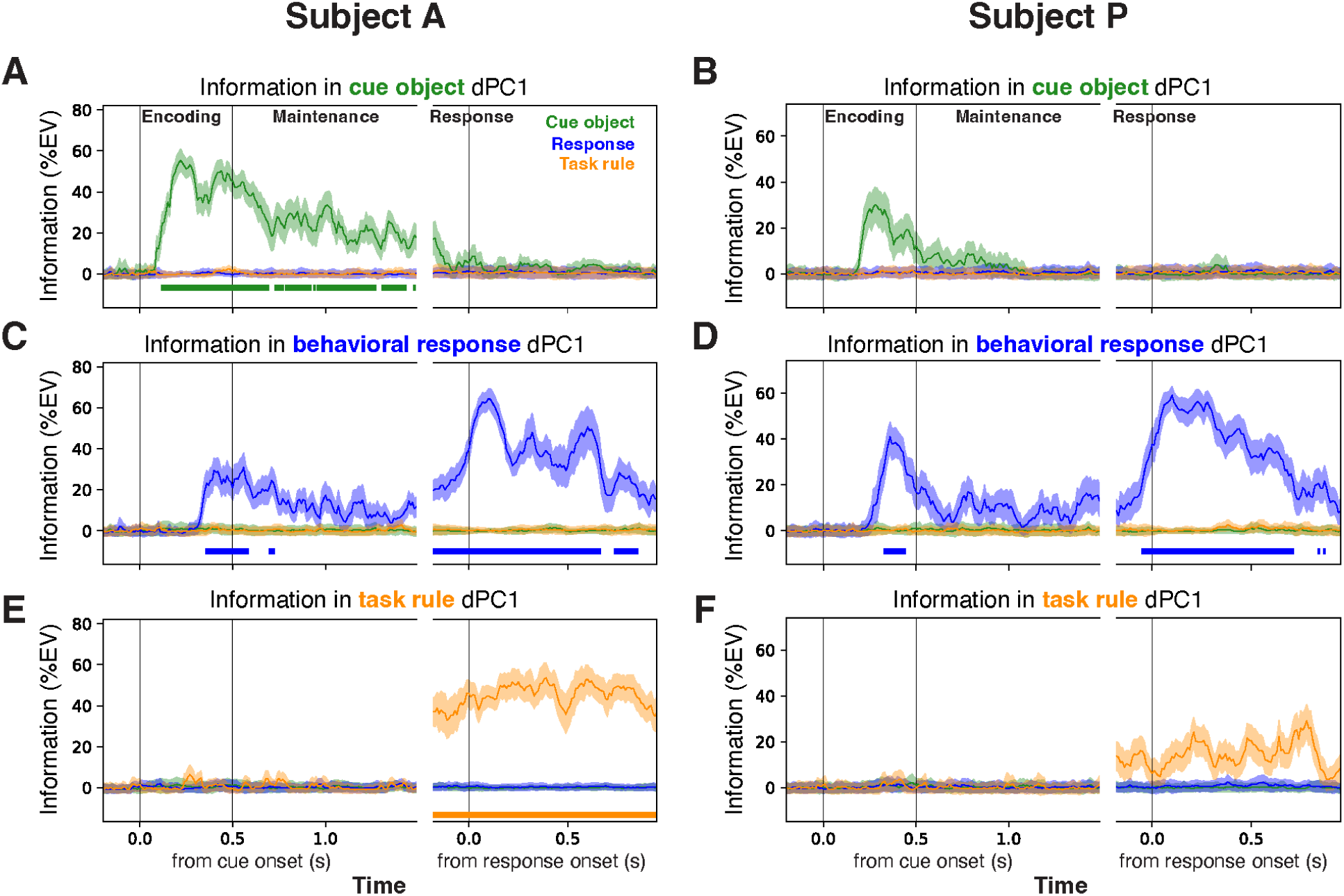
PFC conveyed information about each demixed task factor in Object task (related to Fig. 2). (A,B) Variance explained by cue object identity (green), behavioral response direction (blue), and object→response task rule (orange) for cross-validated PFC population data projected into the dPCA subspace optimized to reflect cue object identity in subject A (A) and subject P (B). Subspace training and testing were performed at the same time points, independently for each time point within the trial. (C,D) Same, but for data projected into the response direction subspace. (E,F) Same, but for data projected into the task rule subspace. For all subspaces, the projected data only conveyed information about the factor the subspace was trained on, indicating dPCA successfully demixed the factors. For the factor each subspace was trained on—analogous to traditional measures of population information^29,30^—PFC activity conveyed information about all factors (significant for all factors in subject A, significant for behavioral response but with similar non-significant trends for other factors in subject P). Information about the cue object appeared at a shorter latency than information about the response direction, consistent with the behavioral response being computed from feedforward inputs reflecting the cue.

**Figure S2.**
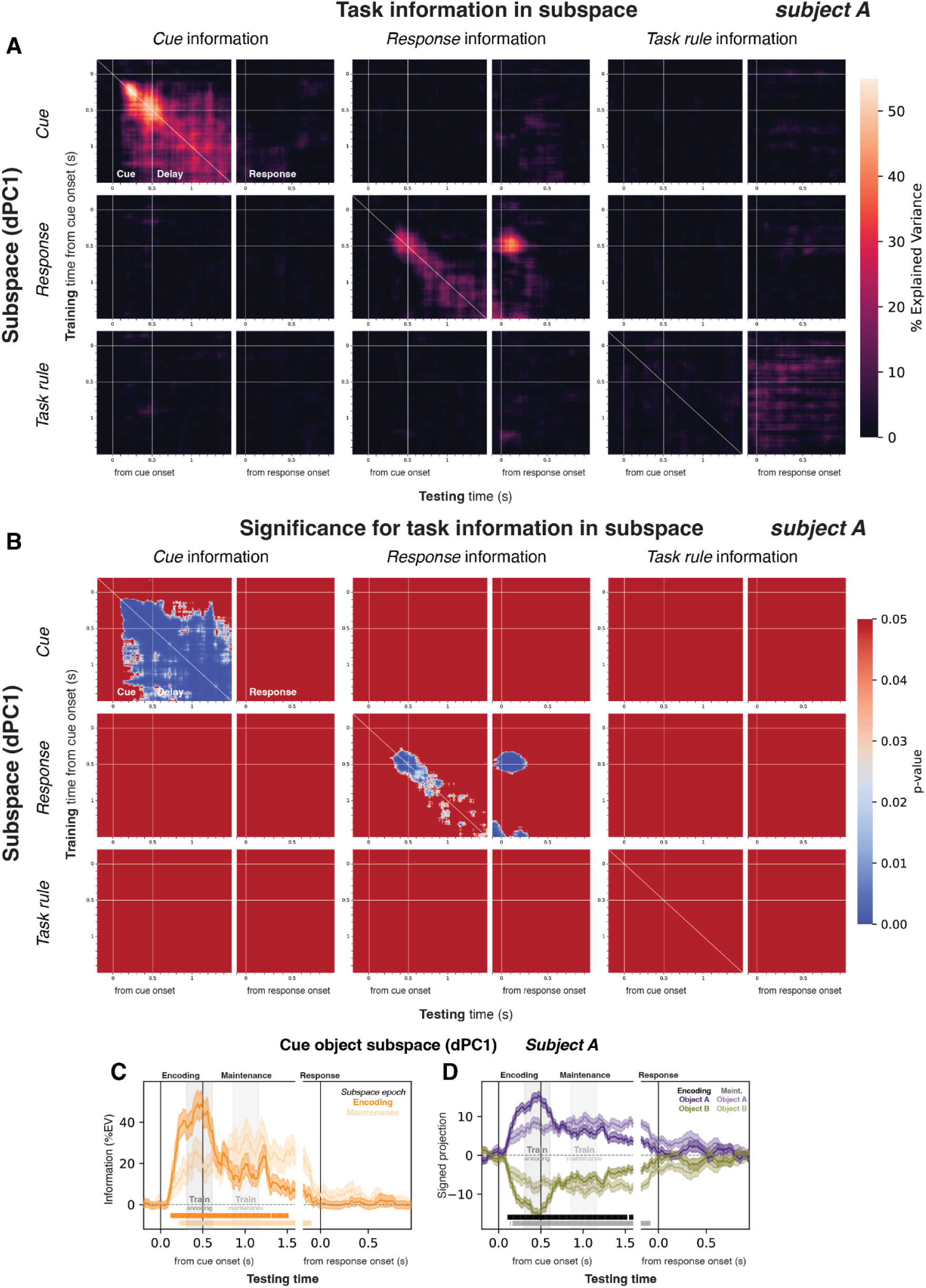
Information for all task variable cross-projections in Object task for subject A (related to Fig. 2). (A) Information (PEV) for all task variable cross-projections. As in Fig. 2, each subplot is a cross-temporal subspace generalization matrix (cf. Fig. 2A), where each value reflects the percent of variance explained (PEV) in data at a given test time point (x-axis), when projected onto a dPCA axis (dPC1) fit at a distinct time point (y-axis). Values along the matrix diagonals reflect data trained and tested at the same time. Off-diagonal values indicate the degree to which a subspace from one time point also conveys information at other times. The full set of subplots corresponds to estimating a dPC axis for each task variable (rows) and measuring PEV of all other task variables (columns) projected onto the fitted first dPC axis (dPC1). Training and testing is performed for all task variables: the cue object identity (top row/left column), the instructed behavioral response direction (middle row and column), and the current task rule giving the mapping from object to response (bottom row/right column). The center subplot is a replotting of Fig. 2B in the main text. There is little information in PFC about the current task rule (lower-right; note that the values plotted in Fig. S1E,F would correspond to an extension of the main diagonal into the response epoch, which is not included here for brevity). In this task, there is little evidence of any cross-generalization between task factors (off-diagonal subplots). (B) Significance for all task variable cross-projections in A. Layout and logic are identical to A. Here, the heatmap values are p values reflecting the significance of the corresponding PEV values. Blue regions reflect regions of significant PEV (bootstrap Z-test, FDR-corrected). (C) Summary of cue object PEV in data projected onto axes fit during encoding (darker curve and fill) and maintenance (lighter) epochs. Cross-projections between the encoding and maintenance subspaces carried information about the cue object that was roughly halfway between their maximal values (full generalization) and zero (no generalization). (D) Mean signed projection onto encoding (darker) and maintenance (lighter) axes for object A (purple) and object B (green) trials. Signs of projections remained consistent throughout the period they were active. These results indicate that, unlike coding of the behavioral response, coding of the cue object was not segregated into fully orthogonal subspaces. Further, there was no evidence of any later subspace reactivation of either cue object subspace.

**Figure S3.**
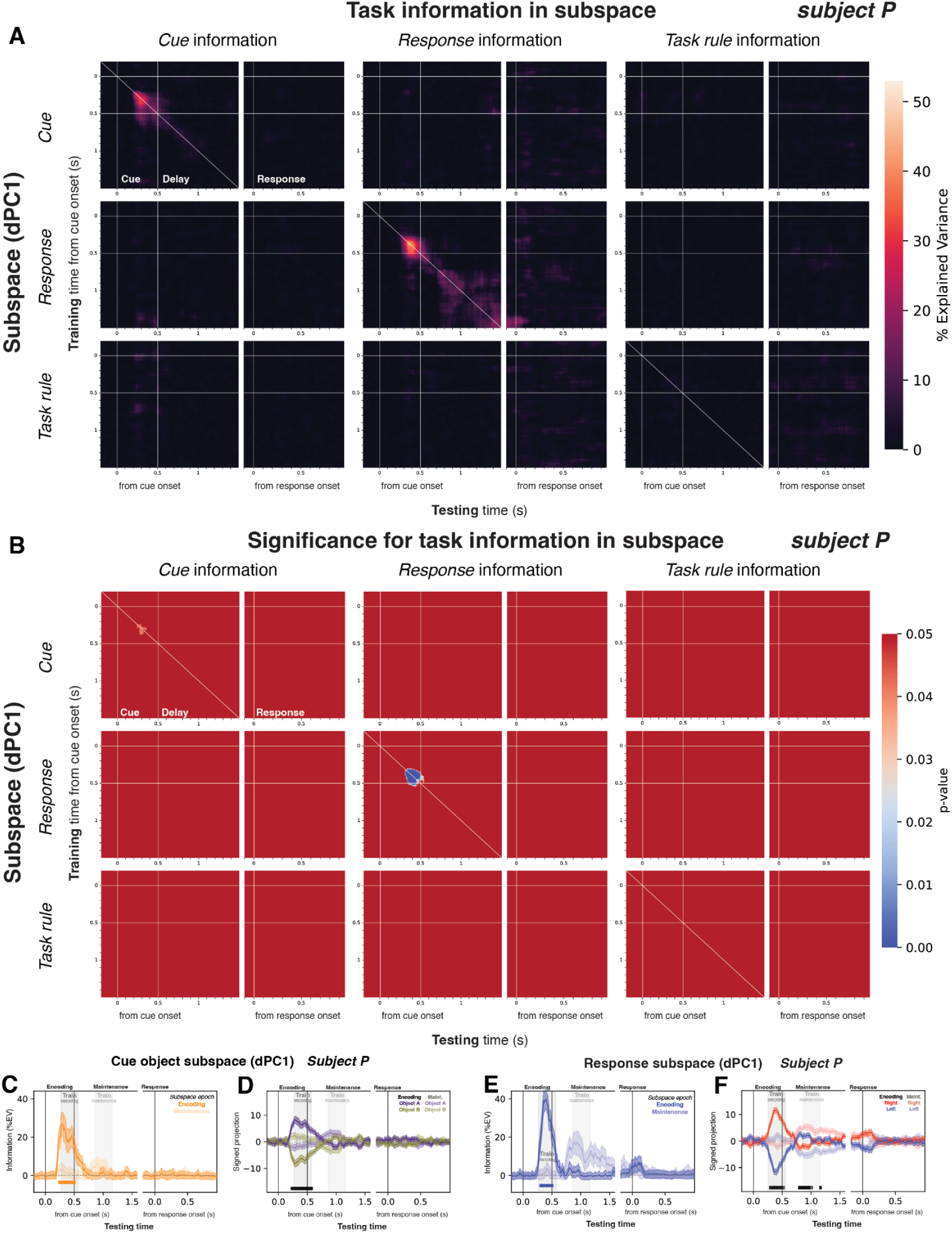
Information for all task variable cross-projections in Object task for subject P (related to Fig. 2). (A) Information (PEV) for all task variable cross-projections for subject P. See Fig. S2 for details. (B) Significance for all task variable cross-projections in A. (C) Summary of cue object PEV in data projected onto axes fit during encoding (darker curve and fill) and maintenance (lighter). (D) Mean signed projection onto encoding (darker) and maintenance (lighter) axes for object A (purple) and object B (green) trials. (E) Summary of response direction PEV in data projected onto axes fit during encoding (darker curve and fill) and maintenance (lighter). (F) Mean signed projection onto encoding (darker) and maintenance (lighter) axes for left (blue) and right (red) response trials. Subject P showed only a non-significant trend toward reactivation of the encoding-epoch subspace during the behavioral response (small bump around “Response” in E and F).

**Figure S4.**
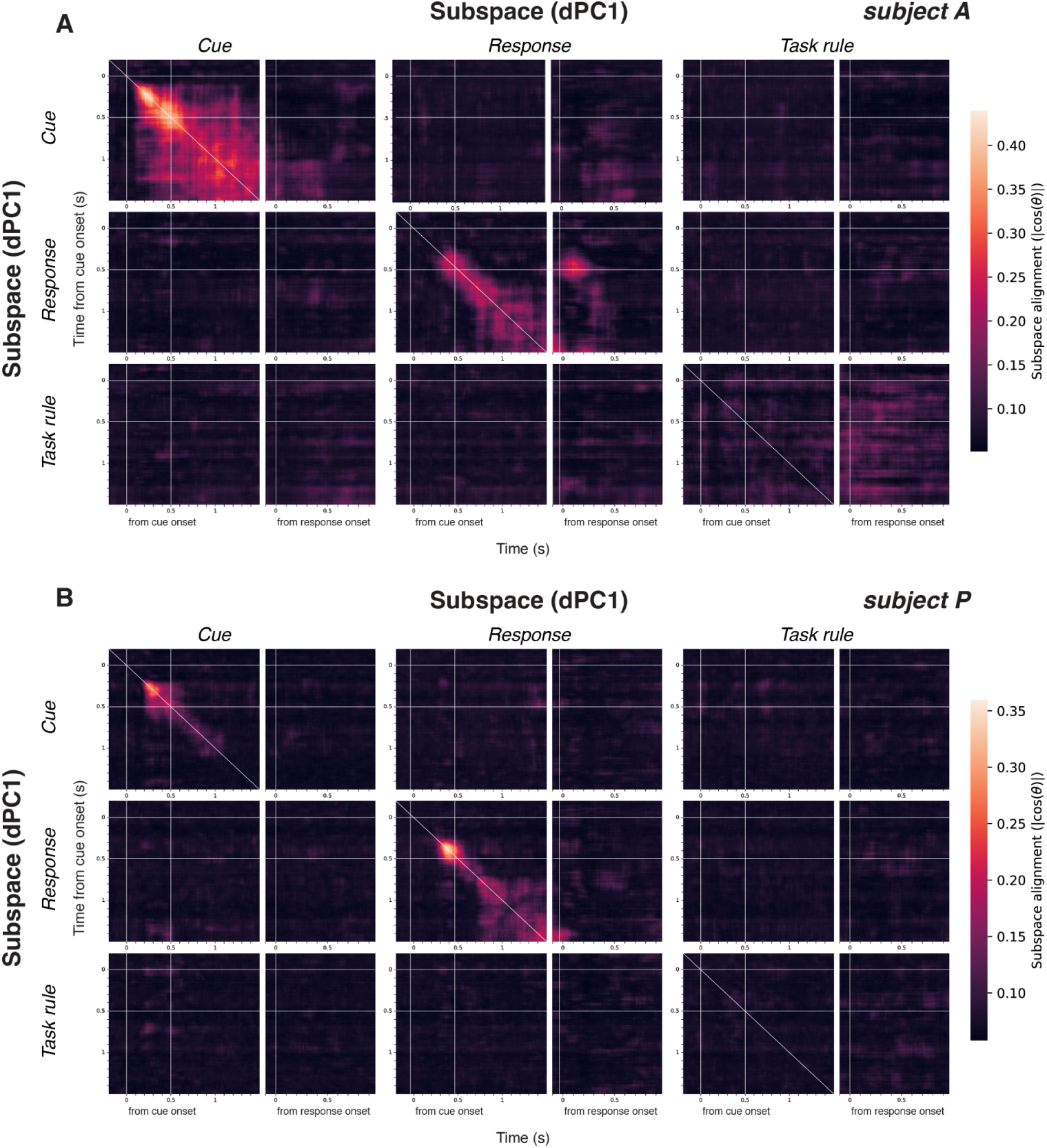
Subspace alignment in Object task (related to Fig. 2). Alignment between subspaces fit at each time point, and for each Object task factor, for subject A (A) and subject P (B). Layout is identical to Fig. S3A, except here the plot matrices are symmetric. Alignment is quantified as the cross-validated absolute value of the angle cosine between the first dPC component for each task factor. It ranges from 0 (indicating compared subspaces are perfectly orthogonal) to 1 (indicating subspaces are perfectly aligned). Points along the main diagonals are the cross-validated alignment of a subspace with itself. Thus, they reflect the maximum expected alignment of that subspace, given the data signal-to-noise ratio. Results indicate that encoding-epoch and maintenance-epoch subspaces for the same task factors are approximately orthogonal (except for the cue object subspace in subject A; upper-left of A). They also show that the reactivation of the encoding-epoch response direction subspace during the behavioral response epoch in subject A (center plot of A) exhibits alignment close to their expected maximal value (points around diagonal).

**Figure S5.**
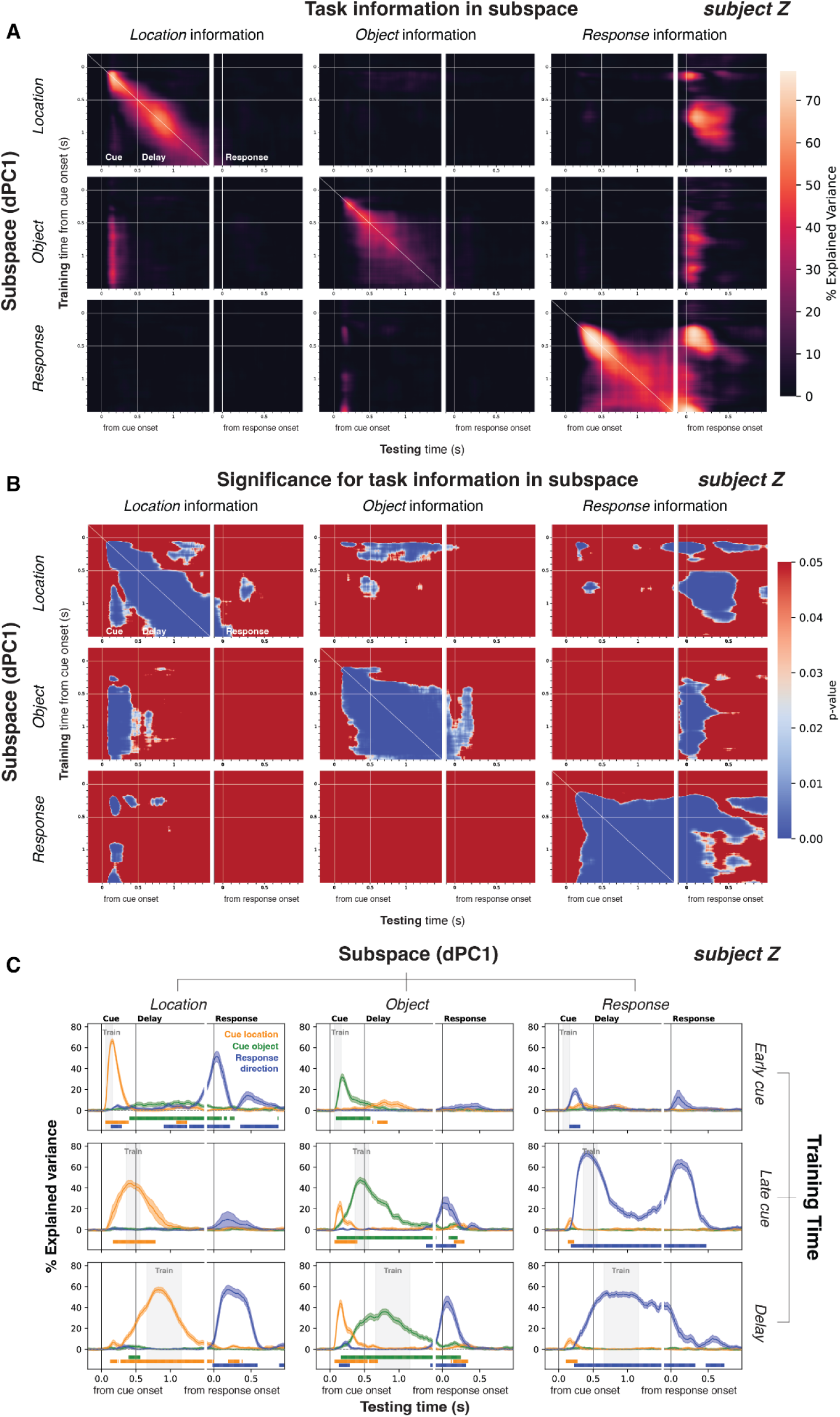
Information for all task variable cross-projections in Object/Location task for subject Z (related to Fig. 3, 4, and 6). (A) Information (PEV) for all task variable cross-projections. The set of subplots corresponds to estimating a dPC axis for each task variable (rows) and measuring PEV of all other task variables (columns) projected onto the fitted first dPC axis (dPC1). Training and testing is performed for all task variables: the cue location (top row/left column), the cue object identity (middle row/center column), and the instructed behavioral response direction (bottom row/right column). The lower-right subplot is a replotting of Fig. 3C in the main text. (B) Significance for all task variable cross-projections in A. As detailed in the main text, the encoding-epoch reponse direction subspace is reactivated during the response (lower-right plots), the encoding-epoch and maintenance-epoch cue location subspaces are recycled to represent response direction during the response (upper-right plots), and the maintenance-epoch cue object subspace is recycled to represent cue location during encoding (middle-left plots) and to represent response direction during the response (middle-right plots). (C) Summary of PEV in data projected onto axes fit to all task factors and trial epochs. Note different subplot layout from A and B. The set of subplots corresponds to estimating a dPC axis for each time epoch (rows) and task factor (columns), and measuring the PEV of all other task variables (curves) projected onto the fitted first dPC axis (dPC1). In each subplot (fitted [time epoch, task factor] pair), PEV is shown for information reflecting the cue location (orange), cue object identity (green), and response direction (blue).

**Figure S6.**
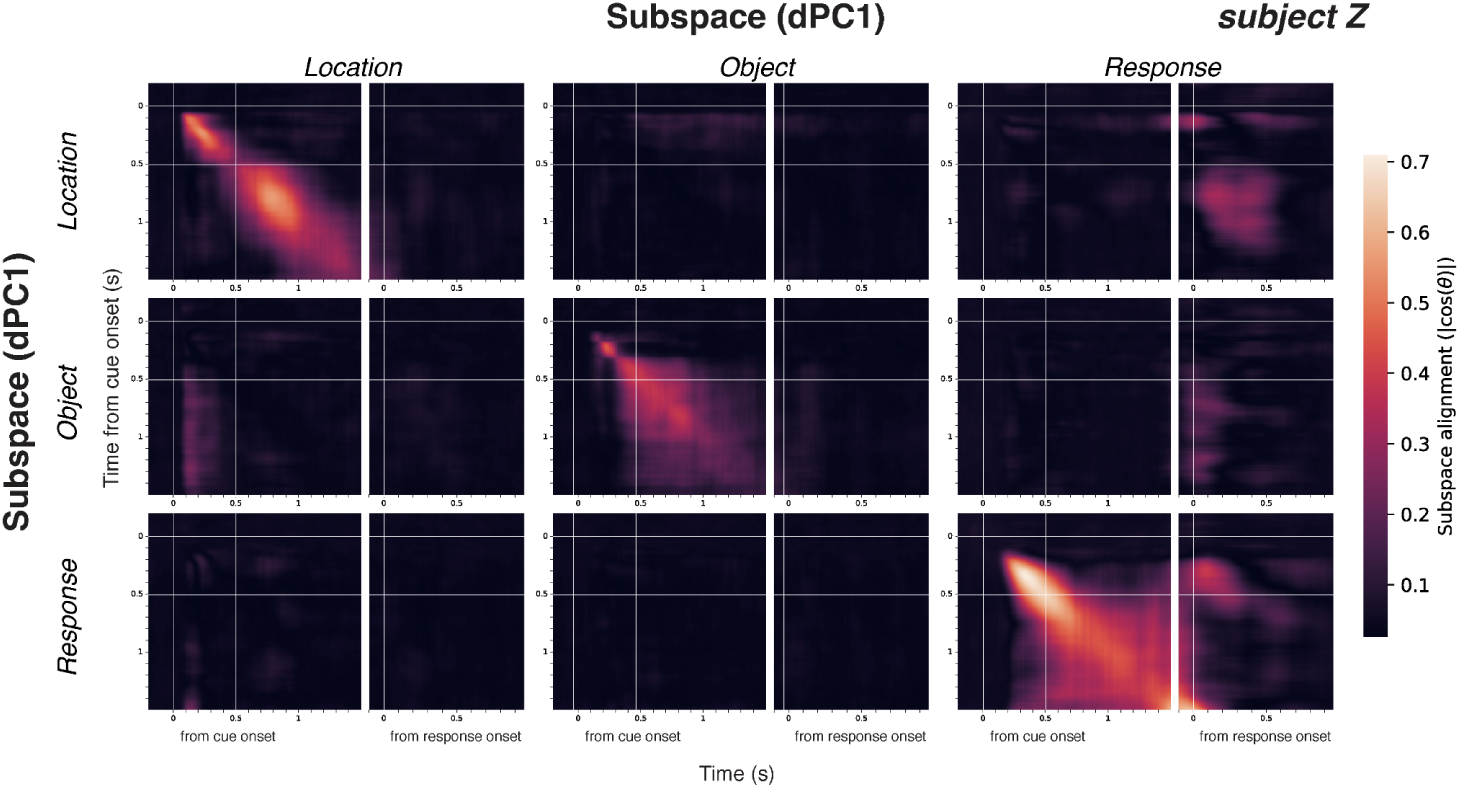
Subspace alignment in Object/Location task (related to Fig. 3, 4, and 6). Alignment between subspaces fit at each time point, and for each Object/Location task factor, for subject Z. Analysis and layout are identical to Fig. S4. Time points around the behavioral response show partial alignment to both the encoding-epoch response direction subspace (lower-right plot) and the encoding-epoch cue location subspace (upper-right plot).

### Supplementary Discussion

We observed somewhat different results between the two datasets analyzed here. In the Object/Location task, subspaces for the cue object (Fig. 6) and cue location (Fig. 4) were later recycled to represent the behavioral response. For the Object task, there was no significant evidence of such subspace recycling (Supplementary Fig. S2,3). One possible explanation for the difference between the two tasks is the level of experience with the learned associations, and thus likely in their associative strength. In the Object/Location task, the data was obtained only after the same familiar associations were well-learned over many months. In the Object task, new associations were learned daily and their reversed mappings were relearned multiple times per session. Our analyses used data just after each reversal relearning episode. Perhaps subspace recycling is observed only in stronger, more well-established representations. Alternatively, the requirement to use both cue identity and location in the Object/Location task could have resulted in “binding” of object and location information together and promoted representational recruitment between them. Under this hypothesis, the lack of any such binding in the Object task prevented recycling of subspaces for different purposes. Distinguishing these hypotheses will require further experimentation.

